# Positive associations between growth and thermal tolerance in the reef-building coral *Montipora capitata*

**DOI:** 10.1101/2025.01.31.635983

**Authors:** Nia S. Walker, Hayley Luke, Spencer Miller, Darienne Kealoha, Carlo Caruso, Erika C. Johnston, Elena M. Mujica, Joshua R. Hancock, Crawford Drury

## Abstract

Relationships between ecologically important traits are increasingly important for the future of coral reefs, which are declining globally due to a variety of stressors. Corals that resist and recover from bleaching will be selected under future climates, which may negatively or positively impact a range of associated traits important for the structure and function of contemporary and future reefs. Using standardized assays, we investigated growth under ambient conditions, bleaching resistance and recovery after bleaching in a model population of 60 *Montipora capitata* colonies harboring diverse symbiont communities. We found significantly higher trait variance within *Durusdinium*-dominated colonies, highlighting the interaction of host and symbiont. We also show that symbiont community impacted thermal tolerance and survivorship during the thermal stress assay, but not recovery or growth. There was also a positive relationship between thermal tolerance and change in surface area and thermal tolerance and recovery from equivalent amounts of stress. These results demonstrate limited tradeoffs to thermal tolerance and suggest that bleaching tolerant corals in this model system are suited to recover from stress and maintain high growth rates under ambient conditions, providing insight into the adaptive capacity of thermally tolerant corals in the Anthropocene.

## Introduction

Although reef-building corals are exposed to a wide variety of stressors which contribute to their decline, ocean warming-induced thermal stress has caused catastrophic coral bleaching and mortality (Hughes et al. 2018, 2017) and is the most certain to persist and intensify (Donner et al. 2018). For example, current models predict that widespread coral bleaching may regularly begin as early as springtime as opposed to late summertime and certain corals may experience year-round bleaching temperatures by 2080 (Mellin et al. 2024). Therefore, thermal stress tolerance, which we have defined in this study as the ability to withstand bleaching and mortality during heating events, is an integral trait for the long-term persistence of coral reefs. Although contemporary stress intensities favor higher thermal tolerance, future climates with more frequent warming events (van Hooidonk et al. 2016) will cause physiological damage to even the most thermally tolerant corals. This forecast means that both basal thermal tolerance and the ability to recover from stress contribute to the long-term persistence of coral reefs (Grottoli et al. 2014, Howells et al. 2021, Hughes et al. 2017). Our coral species of interest, *Montipora capitata*, exhibits strong intraspecific variation in thermal tolerance and bleaching recovery (Cunning et al. 2016, Matsuda et al. 2020, Rodrigues & Grottoli 2007), which makes *M. capitata* an ideal study system for thermal stress resilience– the combined capacity for thermal stress tolerance and recovery.

Thermal tolerance is a complex trait that is significantly influenced by coral host genetics (Barshis et al. 2013, Howells et al. 2021, Thomas & Palumbi 2017) and symbiosis dynamics (Cunning & Baker 2020, Palacio-Castro et al. 2023). In our study system, bleaching during natural warming events is strongly (but not absolutely) associated with divergent symbiont communities (family Symbiodiniaceae), where thermally tolerant corals harbor *Durusdinium trenchii* and *Durusdinium glynnii*, which are traditionally considered to be more stress tolerant (LaJeunesse et al. 2010, Manzello et al. 2019, Palacio-Castro et al. 2023). Conversely, more thermally sensitive corals typically harbor less tolerant symbionts in the diverse genus *Cladocopium,* which also tends to support higher coral growth rates under ambient conditions, representing a tradeoff between growth and thermal tolerance (Jones & Berkelmans 2010), although this effect may be lost under warmer conditions. These findings highlight the importance of symbiont composition in determining thermal stress outcomes, particularly as different symbiont genera are linked to trade-offs between stress tolerance and growth. Focusing on the role of symbionts is therefore crucial to understanding coral responses to climate-induced thermal stress.

Coral thermal tolerance is also linked to other fitness traits, largely because there is an ultimate energetic budget for any organism that must be partitioned into growth, reproduction, resistance to stress and resilience after stress. For example, corals that resist bleaching longer than others have a higher likelihood of post-thermal stress survival, even if they also bleach (Walker et al. 2022, 2023), but tolerant individuals may also grow significantly more slowly than less tolerant corals during bleaching recovery (Walker et al. 2023). These patterns may also reflect interspecific variation, where thermally tolerant species may experience higher mortality during bleaching recovery compared to thermally susceptible species (Matsuda et al. 2020), which bleach and recover. Bleaching also compromises reproductive capacity, where bleaching-resistant corals have higher reproductive output than conspecifics that bleached and recovered (Fisch et al. 2019, Johnston et al. 2020, Leinbach et al. 2021). Some studies have reported tradeoffs between thermal tolerance and growth rates (Bay & Palumbi 2017, Cornwell et al. 2021, Walker et al. 2023), but others have demonstrated no significant relationship (Morikawa & Palumbi 2019). Conversely, there are several instances of positive relationships between thermal tolerance and growth (Lachs et al. 2023, Smith et al. 2007, 2008) and positive associations between other traits such as response to thermal stress and ocean acidification (Wright et al. 2019). Broadly, these findings highlight that coral thermal tolerance exhibits complex dynamics with other fitness traits that may vary within and across species.

In this study, we investigated the complex interplay between growth and thermal tolerance in the reef-building coral *Montipora capitata*. Using 60 adult colonies from a newly established model population, we evaluated symbiont community composition, baseline growth, thermal stress tolerance, and recovery from thermal stress. To ensure robust comparability, we standardized bleaching intensity by exposing corals to the number of days required to reach a bleaching target, allowing us to evaluate differences in the other parameters under standardized thermal stress conditions. Specifically, we explored (i) how Symbiodiniaceae community influenced thermal resilience and growth, (ii) links between thermal tolerance and recovery, (iii) and tradeoffs associated with growth and thermal tolerance. This experimental design provides valuable insights into the multifaceted drivers of coral resilience and the tradeoffs that may shape their response to climate change.

## Materials and Methods

### Sample Collection and Maintenance

We established a model population of 60 *Montipora capitata* colonies at ∼1-2 m depth in a lagoon on Moku o Lo⍰e (Hawai⍰i Institute of Marine Biology–HIMB) in Kāne⍰ohe Bay, O⍰ahu, Hawai⍰i (21°25’57" N 157°47’22" W). We collected at least four ∼5 cm fragments from each colony on August 16, 2022, mounted them on aragonite plugs, and allowed them to recover from sampling stress for 10 days in flowthrough tanks receiving ambient seawater. We recorded temperature with independent loggers in each tank for the duration of acclimation, growout, thermal stress and recovery (Onset U22; Fig. S1). The tank temperatures during the growout and recovery phases were not manipulated and reflect the surface temperature of the lagoon where our source coral population resides. We also recorded light in each tank during thermal stress and recovery, but we did not acquire light data on the acclimation or growout periods; we were able to record light for three weeks following the growout period (November 23 to December 13, 2022) (Odyssey PAR Logger; Fig. S1).

### Growth Experiment

We split replicate ramets of all genotypes across 3 tanks and monitored growth for 12 weeks using buoyant weight and structure-from-motion photogrammetry between August 26 (T_0_) and November 17, 2022 (T_F_). We recorded buoyant weight in triplicate using a bottom-weighing scale while each fragment was fully submerged in seawater. We measured surface area for each ramet using a custom photogrammetry setup with high precision and accuracy (Miller et al. 2024). Briefly, we took 210 photos per coral fragment using a custom 7-camera turntable which creates high overlap and coverage. The setup included 6 Metashape Ground Control Point markers, of which a minimum of 2 were visible in every photo, to allow for standardized measurements across time. We processed photos in Metashape photogrammetry software (version 1.8.4) to build 3D models, trimmed coral fragment 3D models to remove aragonite plugs and collected measurements using AddTools for Metashape add-on software. We calculated percent change of surface area and buoyant weight to quantify skeletal change, and log-transformed data for downstream statistical analyses (Fig. S2).

### Evaluation of Symbiodiniaceae Communities

We collected three additional fragments (∼2 cm^2^) from each colony on September 29, 2022 and preserved them in 70% ethanol. We extracted genomic DNA using the OMEGA E-Z 96 Tissue DNA Kit, following manufacturer protocol (Bio-Tek). We targeted the ribosomal internal transcribed spacer 2 (ITS2) region using the ITS-DINO (TCG TCG GCA GCG TCA GAT GTG TAT AAG AGA CAG GTG AAT TGC AGA ACT CCG TG) (Pochon et al. 2001) and ITS2Rev2 (GTC TCG TGG GCT CGG AGA TGT GTA TAA GAG ACA GCC TCC GCT TAC TTA TAT GCT T) (Stat et al. 2009) primers, which were modified to include Illumina forward and reverse sequences. We conducted PCR amplification with the following protocol: initial denaturation at 95°C for 3 min; 30 cycles of 95°C for 30 sec, 55°C for 30 sec, and 72°C for 30 sec; and a final extension at 72°C for 5 min. We cleaned ITS2 amplified DNA using Mag-Bind TotalPure NGS beads, following manufacturer protocol (OMEGA Bio-Tek). We conducted a second round of PCR to attach unique dual Nextera XT primers (Set A and D) to individual samples; the thermal profile was the same as above except for utilizing 25 PCR cycles. Both PCR rounds used Hotstart 2x mastermix. We cleaned libraries using Mag-Bind TotalPure NGS beads, normalized after determining concentrations (QuantIT kit, Qiagen), and pooled for sequencing on the Illumina MiSeq v3 platform with 2×300 paired end reads (University of Hawai‘i at Mānoa Advanced Studies in Genomics, Proteomics and Bioinformatics). We used SymPortal (Hume et al. 2019) to generate ITS2 type profiles that include genus, species, and population level designations for *Symbiodiniaceae* from submitted fastq files.

We retrieved high-quality reads for ITS2 identification from the 180 samples, representing all 60 genotypes in triplicate. Samples initially yielded 8,148 to 459,327 ITS2 reads, on average 35,006 reads. Following SymPortal quality filtering, sample reads ranged from 2,335 to 349,971 reads, averaging 21,851 reads. We determined genet level type profiles by consolidating all ramet type profiles. We then categorized symbiont classifications into *Cladocopium*-dominated (≥ 80% *Cladocopium* reads), *Durusdinium*-dominated (≥ 80% *Durusdinium* reads), or Mixed (80% > *Cladocopium* reads > 20%).

### Thermal Performance Assay

We conducted the thermal performance assay from January 23 to February 5, 2023. Coral fragments from each genotype were divided across 3 replicated heated tanks and one control tank. The 14-day thermal stress profile (which resolves naturally occurring bleaching phenotypes; Drury et al. 2022) followed a nonlinear ramp from 25°C to 32°C over the first 5 days and was maintained at 32°C for the remaining 9 days (Fig. 1C). The control tank remained at ambient temperature (∼25°C) throughout. Each morning (10:00AM), we scored ramets as alive or dead based on the presence of any living coral tissue.

**Figure 1:**
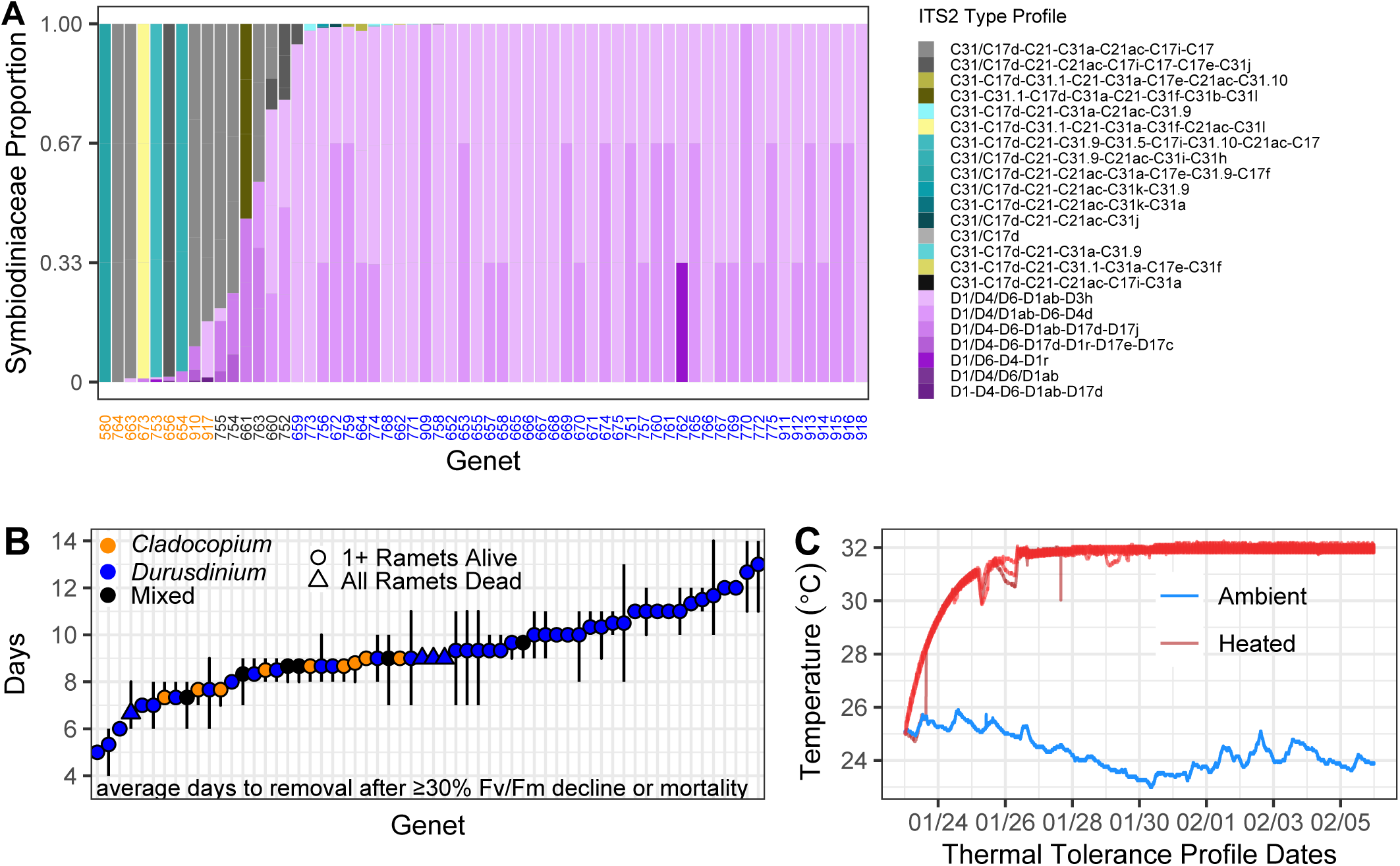
Symbiodiniaceae composition and temperature ramp design. (A) Barplot showing relative proportions of SymPortal Symbiodiniaceae type profiles by genotype across 3 replicate fragments. Genet x-axis labels represent genus level symbiont designations. Orange = *Cladocopium*-dominant (≥ 80% *Cladocopium*); Black = Mixed (*Cladocopium* 20-80%); and Blue = *Durusdinium*-dominant (≥ 80% Durusdinium). (B) Average number of days per genet to reach F_v_/F_m_ ≥30% decline, including lines to show the raw spread of ramet variation. Genets with at least one surviving ramet are represented by circles. There were four genets (triangles) where all ramets died before bleaching, and the average number of days to mortality is shown. Color reflects the dominant symbiont community. (C) Thermal stress temperature records, recorded at 1 min intervals. All heated tanks are in shades of red (3 adult and 3 juvenile tanks), and the one adult control tank is in blue.

We characterized symbiont photosynthetic efficiency (maximum quantum yield; F_v_/F_m_) before the experiment and daily throughout the heat-stress assay using pulse-amplitude modulated fluorometry (Walz MAXI Imaging PAM), following a 2-hour dark adaptation period from 4:00 to 6:00PM. We measured F_v_/F_m_ in triplicate for each ramet and removed ramets upon reaching ≥ 30% decline in F_v_/F_m_ compared to pre-thermal stress F_v_/F_m_, exposing corals to a variable number of heat exposure days to achieve similar levels of bleaching. We chose a ≥ 30% decline in F_v_/F_m_ to elicit a visible bleaching response and increase survival for use in the thermal stress recovery experiment.

### Thermal Stress Recovery Assay

We moved bleached fragments into ambient temperature recovery tanks (∼25°C) after reaching a ∼30% F_v_/F_m_ decline from pre-thermal stress F_v_/F_m_. We assessed mortality daily for one week, then weekly up to one month, after two months, and lastly after 80 days of recovery, when we also remeasured F_v_/F_m_.

### Statistical Analyses

We conducted all analyses in R (version 4.3.1). We calculated experimental degree heating weeks (eDHW) from the thermal stress ramp profile (Drury et al. 2022, Leggat et al. 2022) using 27.4⍰ as the mean monthly maximum temperature (MMM). We used a Weibull 1.3 model in the R package *drc* to model relative F_v_/F_m_ for all replicate fragments of each genet against eDHW and calculated the effective dose (in eDHW) required to reach a 30% F_v_/F_m_ decline per genet following Drury et al. 2022. We calculated ED30 because it was the maximum photosynthetic decline in our experimental design. Higher ED30 values represent more accumulated thermal stress required to produce the same degree of physiological decline in each coral genotype. There was one genet (765) for which we could not properly fit a dose response curve, which was excluded from downstream thermal tolerance analyses (Fig. S3).

We used a Kaplan Meier survivorship analysis to examine mortality during the recovery phase, comparing control and heated corals, but did not generate survivorship curves for thermal tolerance because we removed ramets from the heat ramp as they reached a predetermined bleaching threshold. We compared thermal tolerance and survival at the end of recovery using a generalized linear model (survival ∼ tolerance score, family = “binomial”). We compared survivorship at the end of the stress test between symbiont communities (*Cladocopium*, Mixed, *Durusdinium*) and at the end of recovery using a Fisher’s exact test.

We calculated percent change in buoyant weight and surface area (final-initial/initial) for each fragment, calculated the log-transformed mean for each genotype, and compared both metrics to the thermal tolerance score (ED30) using a linear model. We used a Levene’s test to compare variances between symbiont groups for surface area, buoyant weight and thermal tolerance. When variances were significantly different between groups (i.e., thermal tolerance), we used Welch’s ANOVA and a post-hoc Gamer-Howell test, which are robust to unequal group variances, to compare traits between symbiont communities. Otherwise, we compared log-transformed surface area and buoyant weight between symbiont communities using a one-way ANOVA and post-hoc Tukey test.

## Results

### Thermal performance and symbiont community

Coral genotypes averaged 5-13 days in the heat treatment before reaching 30% decline in F_v_/F_m_. Within each genotype, responses were highly consistent and all ramets reached the removal threshold within 2 days of each other in 46 out of 60 genotypes (Fig. 1B, Fig. 2A). ED30 values averaged 4.65 ± 1.14 (mean ± 1SD), with a range of 1.91 to 7.06 eDHW.

**Figure 2:**
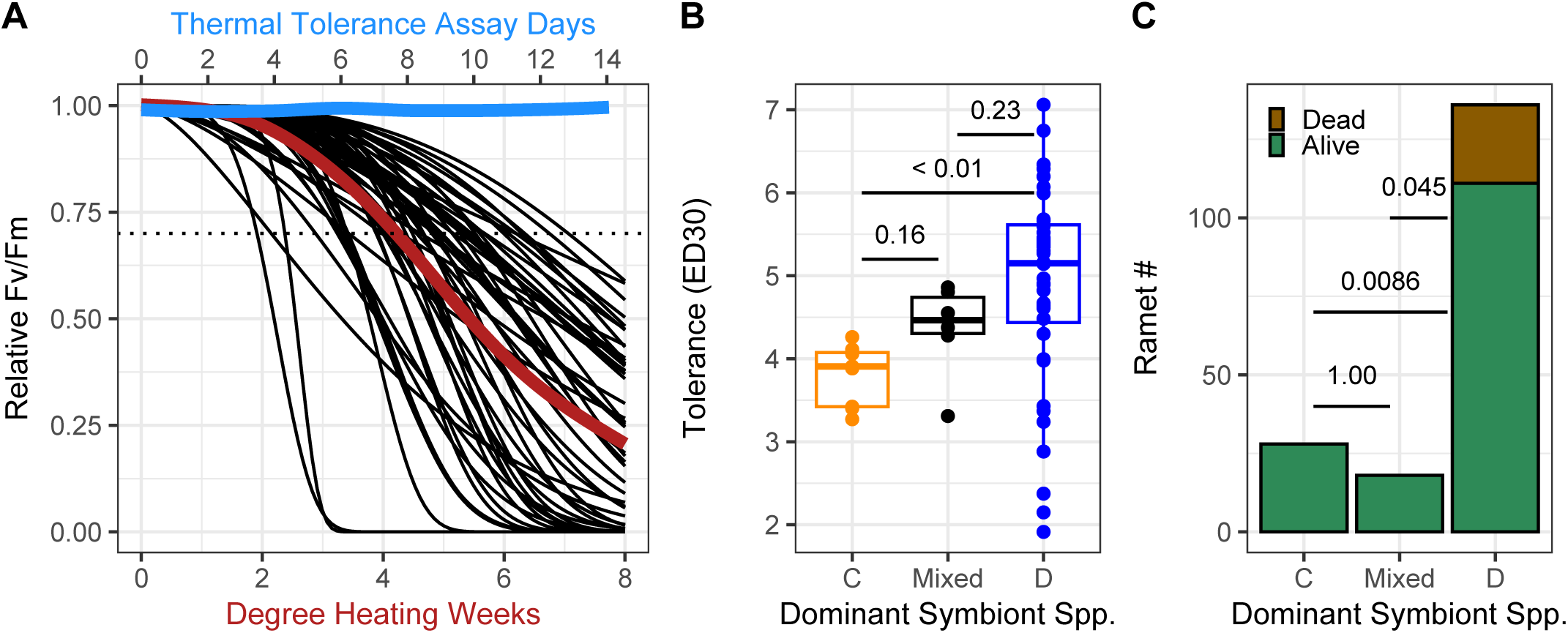
Relationship between thermal tolerance and symbiont community. (A) Summary of dose response curves based on relative F_v_/F_m_ and degree heating week data. The red line represents the average ED30 curve among all heated samples, and the blue line represents average relative F_v_/F_m_ data among controls. The dotted line represents 70% F_v_/F_m_ relative to pre-thermal stress, which is used to calculate the ED30 value (heat dosage required to elicit a 30% decline in photosynthetic efficiency). The bottom x-axis shows degree heating weeks expected to induce F_v_/F_m_ decline in heated corals. The top x-axis refers to the 14-day thermal tolerance assay period, during which non-heated controls were also measured. Each black line is a coral genotype. (B) Thermal tolerance compared to symbiont community, evaluated using a one-way Welch’s ANOVA and post-hoc Gamer-Howell test. Thermal tolerance is based on ED30 curves. (C) Ramet mortality at the end of the thermal tolerance assay by symbiont community, evaluated using a fisher’s exact test.

SymPortal identified 32 Symbiodiniaceae types, most of which were *Cladocopium* (n = 21, 65.6%). These profiles were consolidated Symbiodiniaceae types into 23 type profiles, of which 16 belonged to *Cladocopium*. Most *Durusdinium* profiles were identified as *Durusdinium glynnii* (D1, D4, or D6) (Wham et al. 2017). Most corals were dominated by *Durusdinium* spp. (n = 45, 75%); there were 9 colonies dominated by *Cladocopium* spp., and 6 with mixed communities (Fig. 1A). Symbiont community significantly impacted thermal tolerance (p=0.0012), where *Durusdinium*-dominated corals exhibited significantly higher thermal tolerance (4.88 ± 1.23 ED30) than *Cladocopium*-dominated corals (3.82 ± 0.362 ED30; Games-Howell post-hoc *p* < 0.01), but not mixed corals (4.36 ± 0.565 ED30; Games-Howell post-hoc *p* = 0.23, Fig. 2B). There was a strong relationship between symbiont type and mortality, where the *Durusdinium*-dominated ramets experienced significantly higher mortality than *Cladocopium*-dominated and mixed corals (Fisher’s exact test, *p* = 0.009 and *p* = 0.045, respectively, Fig. 2C).

### Recovery Assay

86.3% (157 out of 182) of ramets survived the thermal stress assay. There were 4 genotypes that experienced total mortality before reaching the bleaching threshold, all of which were *Durusdinium*-dominated (Fig. 1B).

Coral fragments were transferred to an ambient recovery tank and monitored for 80 days after reaching the bleaching threshold; at least one ramet from 29 of 60 genotypes survived the entire recovery period. Within 7 days, we observed 61.1% mortality of ramets in the recovery assay and after 80 days of recovery, mortality increased to 68.2% (Fig. S4A). Thermally stressed corals had significantly lower survival probability than controls (Kaplan Meier log-rank test, *p* < 0.001), with predicted cumulative survival dropping below 50% at one-month after thermal stress (Fig. 3A). All surviving ramets but one (753C) recovered to within 30% of pre-thermal stress F_v_/F_m_ by the end of recovery (Fig. 3B). There was no significant difference in survivorship at the end of the recovery period between corals harboring different symbiont communities (Fig. S4B). There was a strong positive relationship between a genotype’s thermal tolerance and likelihood of ramet survival during the recovery period (logistic regression, *p* = 0.001, Fig. 3C), where more tolerant corals exhibited higher survival, despite experiencing the same physiological damage during the stress.

**Figure 3:**
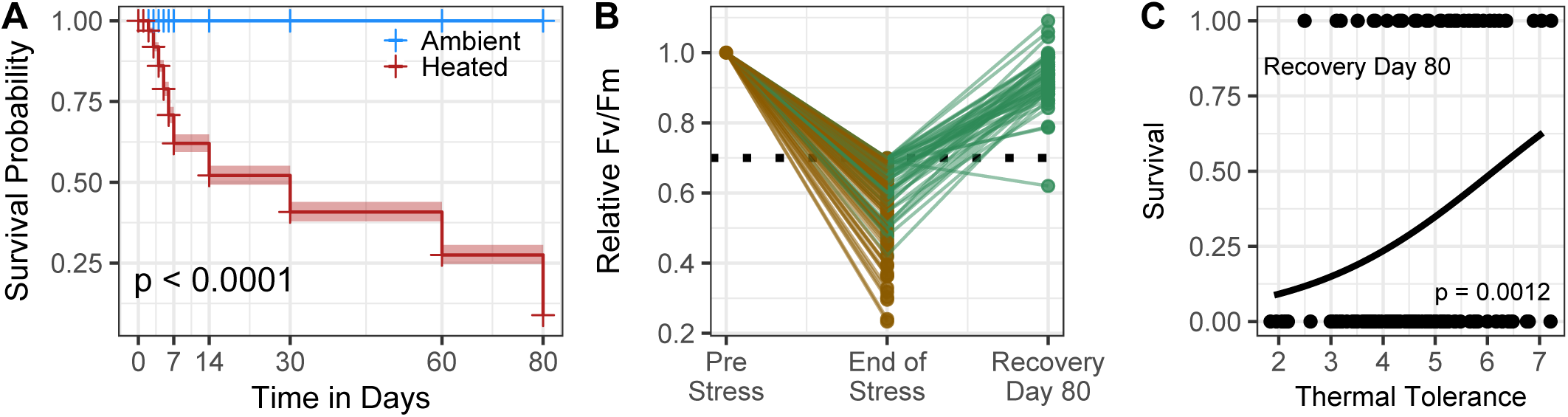
Adult thermal stress recovery. (A) Kaplan Meier survival probability in each treatment across the 80-day post-thermal stress experiment period. (B) Heated ramet relative F_v_/F_m_ upon removal from the thermal tolerance assay and after 80 days of recovery, compared to pre-thermal stress. Green represents ramets that survived to recovery day 80. Brown represents ramets that died between thermal stress removal and recovery day 80. The dotted line denotes 70% F_v_/F_m_ relative to pre-thermal stress. (C) Ramet mortality at the end of recovery compared to the genotype’s average thermal tolerance. The black line represents the logistic regression model best fit. Only ramets that were alive on Day 0 of the recovery period were included in this analysis.

### Tradeoffs with thermal tolerance

During a three-month period prior to thermal stress, all genotypes increased in surface area (range: 1.30% to 19.8%) and buoyant weight (range: 11.8% to 43.6%) (Fig. S2). There was a significant positive linear relationship between thermal tolerance and change in surface area (*p* = 0.0025, R^2^ = 0.041; Fig. 4A). There was no significant linear relationship between thermal tolerance and change in buoyant weight (*p* = 0.93 and R^2^ = 0.0001; Fig. 4C). Although the most thermally tolerant corals grew more than the least tolerant corals, moderately tolerant corals exhibited the highest growth. While coral genotype significantly impacted change in surface area and buoyant weight (Fig. S2, *p* < 0.001), there was no significant relationship between symbiont community and change in surface area (p=0.581, Fig. 4B) or change in buoyant weight (p=0.907, Fig. 4D).

**Figure 4:**
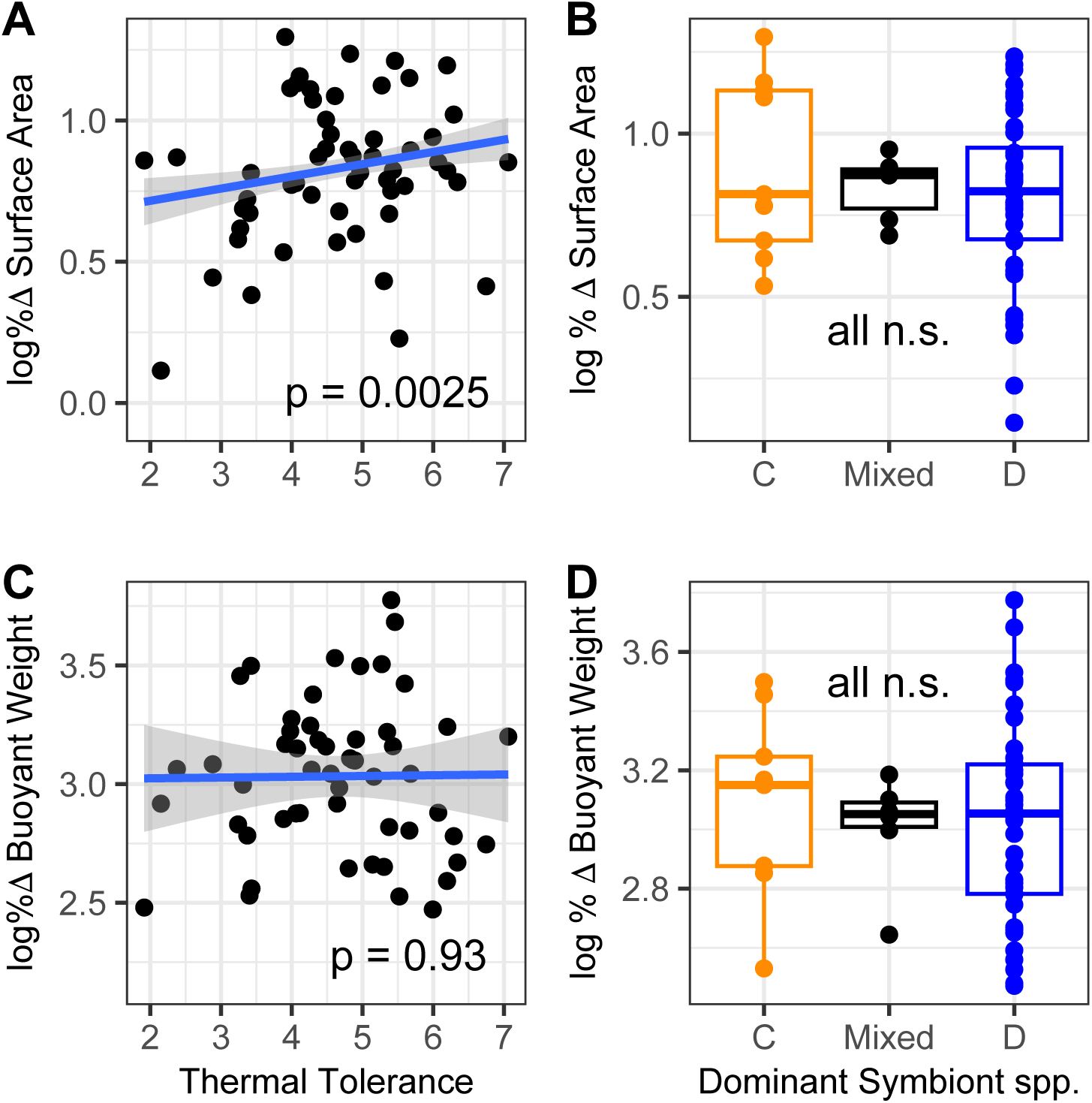
Relationships between thermal tolerance and growth. (A,C) Log-transformed percent change surface area (A) or buoyant weight (C) compared to average thermal tolerance of each genotype. The blue line represents the linear model best fit. (B,D) Log-transformed percent change surface area (B) or buoyant weight (D) compared to the dominant symbiont community of each genotype, each evaluated using a one-way ANOVA and post-hoc Tukey test but all comparisons were not significant.

## Discussion

As coral reefs are increasingly exposed to ocean warming (Hughes et al. 2018, 2017), it is important to understand the relationships between thermal stress tolerance, recovery, and other fitness traits. Here we coupled a growth assay under ambient conditions with a standardized thermal stress assay which exposed corals to similar levels of physiological damage across a range of cumulative thermal stress (Walker et al. 2023) to evaluate the relationship between thermal tolerance, recovery, growth and symbiont dynamics in the reef-building coral *Montipora capitata*.

Our model population was significantly skewed toward *Durusdinium*-dominated corals, likely due to the shallow depth of our collection site. *Durusdinium* is a high-light specialist and is expected to be the dominant community in approximately 70% of *M. capitata* colonies at depths <1m in Kāne⍰ohe Bay (Innis et al. 2018), which closely matches our results (75%). *Cladocopium* species may confer energetic advantages that result in higher prevalence of *Cladocopium*-dominated corals in deeper areas, whereas *Durusdinium* corals may be better adapted to shallow or variable thermal environments (Goulet et al. 2019, Innis et al. 2018, Terraneo et al. 2023).

*Durusdinium*-dominated corals exhibited significantly higher thermal tolerance than *Cladocopium*-dominated corals, reproducing the canonical tolerance patterns of these genera and mirroring responses during natural bleaching events in Kāne⍰ohe Bay (Cunning et al. 2016). Although our sample size of *Durusdinium-*dominated corals is substantially larger than *Cladocopium*, we also found these corals had significantly higher variance in thermal tolerance and that the least tolerant corals in the population also harbored *Durusdinium*. In our experiment, all *Cladocopium*-dominated corals reached the F_v_/F_m_ decline threshold within 9 days, while most *Durusdinium*-dominated corals retained their symbiotic efficiency (60% exceeded 9 days) at the risk of sustaining lethal levels of thermal stress. We only observed mortality during the stress-test in these *Durusdinium*-dominated corals, which may indicate a mismatch between stable function of the symbiont community and extreme stress in the coral host. Overall, this pattern and the higher variance in *Durusdinium-*dominated corals reflects the complex interactions between host and symbiont which govern these traits and correspond to both genetic and symbiotic drivers of thermal tolerance in this species.

Previous work has demonstrated patterns of lower mortality in *Durusdinium* compared to *Cladocopium*-dominated corals (Cunning et al. 2016, Dilworth et al. 2021, Turnham et al. 2023) by testing corals that experienced the same amount of heat exposure. Conversely, we present a functional thermal stress assay in which corals were exposed to different lengths of heat to reach the same F_v_/F_m_ decline. Our findings point to a decoupling of the coral-symbiont symbiosis under varying amounts of thermal stress time depending on the symbiont type.

*Durusdinium*-dominated colonies typically have higher symbiont densities but lower photopigments per symbiont cell and translocate less carbon to the host in this system (Wall et al. 2020), both of which interact with the light environment of the coral. These traits would be expected to compromise growth, and previous work dividing *M. capitata* into binary symbiont groups demonstrated divergent baseline growth rates between genera; where cooler months favored *Cladocopium*-dominated corals, but warmer (still non-stressful) months elicited no difference in growth between symbionts (Matsuda et al. 2023). We found no relationship between these factors when coral growth was measured around 27.5⍰ (median during growth assay, August to November). This warmer temperature relative to Matsuda et al. (2023) may have eliminated or reduced the benefit of *Cladocopium* and corresponds to a loss of growth tradeoff at warmer temperatures previously described in *Pocillopora* (Cunning et al. 2015). Our growth assay was also 3 months and designed to capture more integrative growth across a season, which may obscure more fine-scale differences based on temperature.

Overall, our results show limited evidence for tradeoffs to thermal tolerance in this model system. We found a positive relationship between change in surface area and thermal tolerance. Interestingly, there was no relationship between change in buoyant weight, a proxy for calcification rate, and thermal tolerance, which could point to preferential investment into volumetric growth at the expense of producing denser skeleton in thermally tolerant corals. We also found a positive relationship between recovery and thermal tolerance when corals were exposed to approximately the same physiological damage across a range of accumulated temperature stress. The most thermally tolerant corals had a significantly higher likelihood of survival during recovery, but this pattern was independent of symbiont community. These results support previous work that documented this pattern (Walker et al. 2023) and suggest a diversity of thermal tolerance phenotypes should be used in restoration actions based on contemporary climates, although this relationship may break down in warmer climates.

Positive relationships between thermal tolerance and other traits imply that the selection of thermally tolerant corals will improve growth in cooler seasons and recovery from thermal stress in this model system. Genetic covariance, where the genomic drivers of one fitness-related trait also relates to other traits, could explain this pattern and would suggest a reinforcing effect of climate change on ambient growth and recovery after bleaching. The latter is especially important because bleaching effectively induces a metabolic breakdown which may lead to coral starvation if the coral host cannot meet its energy requirements through increased heterotrophy (Grottoli et al. 2006); corals can clearly recover from this state, but temperature anomalies that are too frequent, persistent or intense may limit the capacity for bleaching recovery. Although positive genetic covariance may increase or decrease the rate of adaptation, a relationship between these two traits could be a particularly important relationship for the long-term persistence of coral reefs.

### Conclusions and Future Directions

This study conducted growth and thermal stress resilience assays on *Montipora capitata* with a range of symbiont communities to evaluate a range of traits. We demonstrate that *Durusdinium*-dominated corals were more tolerant than those dominated by *Cladocopium* spp. and there is a positive relationship between thermal tolerance and both growth and recovery. These results provide valuable insights into putative tradeoffs associated with coral thermal tolerance and growth and links between thermal tolerance and recovery. This study employed a relatively short-term artificial thermal stress assay (McLachlan et al. 2020) and could also benefit from evaluation of traits associated with resilience on longer timescales and under environmental conditions expected in future oceans.

## Supporting information

Onset U22

downstream statistical analyses

thermal tolerance analyses

mortality increased to 68.2

## Acknowledgements

This research was conducted in Kāne⍰ohe Bay, O⍰ahu, Hawai⍰i. We appreciate the enduring significance of coral reefs in traditional Hawaiian culture (Gregg et al. 2015) and hope to honor the relationship between Hawaiian culture and the sea with our research. We also thank Hanalei Ho⍰opai-Sylva, Shayle Matsuda, Khalil Smith, Amro Zayed, Clarisse Sullivan, Giada Tortorelli, Sarah Woo, Kira Hughes, and the Coral Resilience Lab for field, experimental and administrative support. This work was funded by the Paul G. Allen Family Foundation and NOAA (NA20NMF4820290). Fragments were collected under Hawai⍰i DLNR permits SAP 2022-22 and SAP 2023-31 to the Hawai⍰i Institute of Marine Biology.

## Author Contributions

NSW and CD conceived the experiment. NSW, HL, CC, ECJ, JRH, EMM, and CD conducted growth, and thermal stress experiments. NSW, HL, DK, and SM collected data. NSW and CD analyzed data. NSW wrote the manuscript and all authors edited and approved the final manuscript.

## Data Availability Statement

The data that support the findings of this study currently reside in a private GitHub repository but can be shared upon request.

## Supplementary Materials Captions

**Figure S1:**
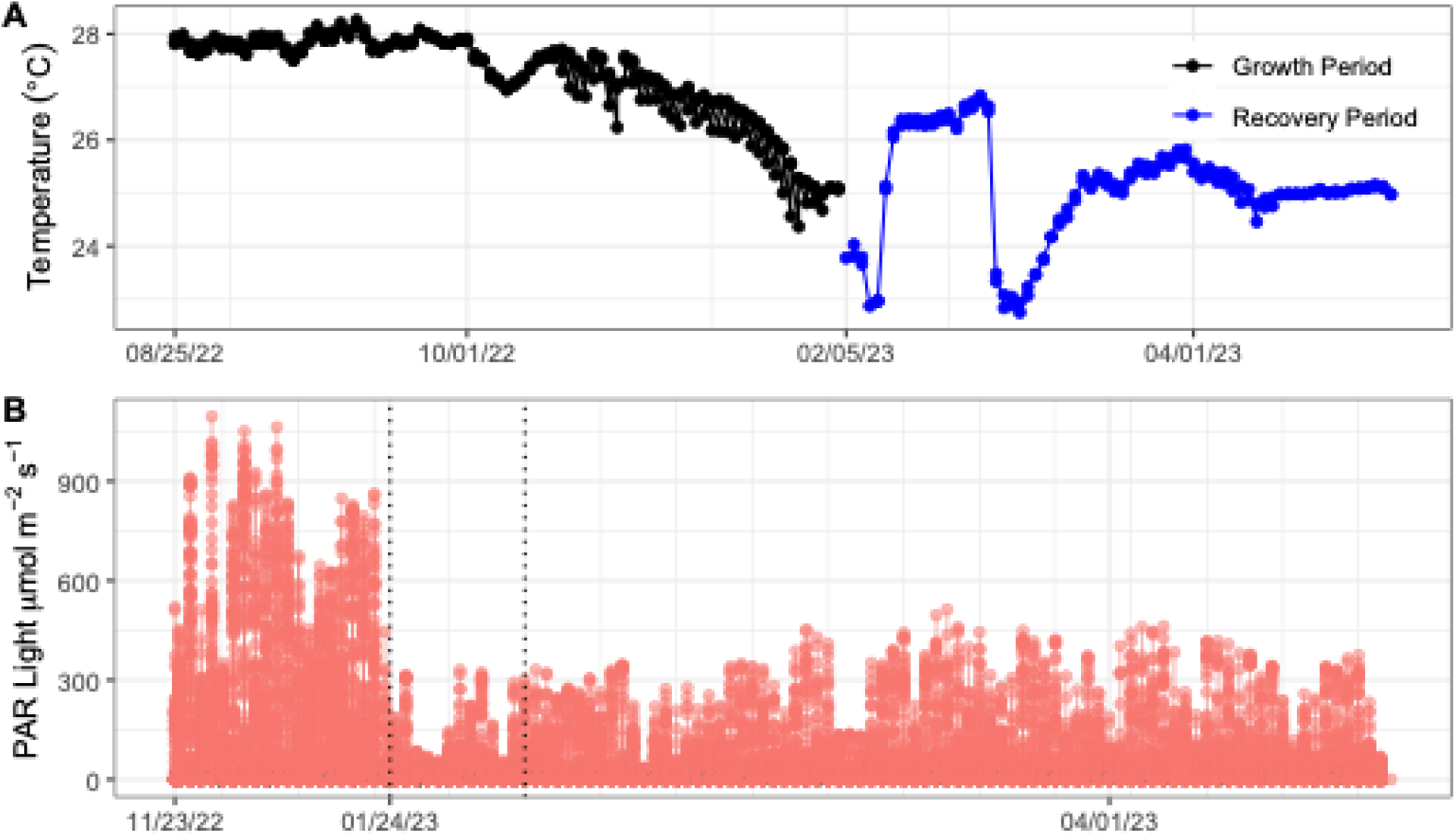
Experimental tank system conditions. (A) Temperature logger profiles of the tanks during the baseline growth period (August 26^th^ to November 17^th^, 2022), and then the recovery period (February 6^th^ to April 26^th^, 2023). All tanks were maintained with ambient seawater. Stress testing temperatures in between growth and recovery are shown in Figure 1C. (B) Light logger profiles for the growout proxy period (11/23-12/13/22, three weeks after the growout period but light data for the growout period 8/26-11/16/22 was unavailable), the thermal tolerance assay (1/23-2/5/23, which is highlighted within two vertical dotted lines), and the thermal recovery period (2/6-4/26/23).

**Figure S2:**
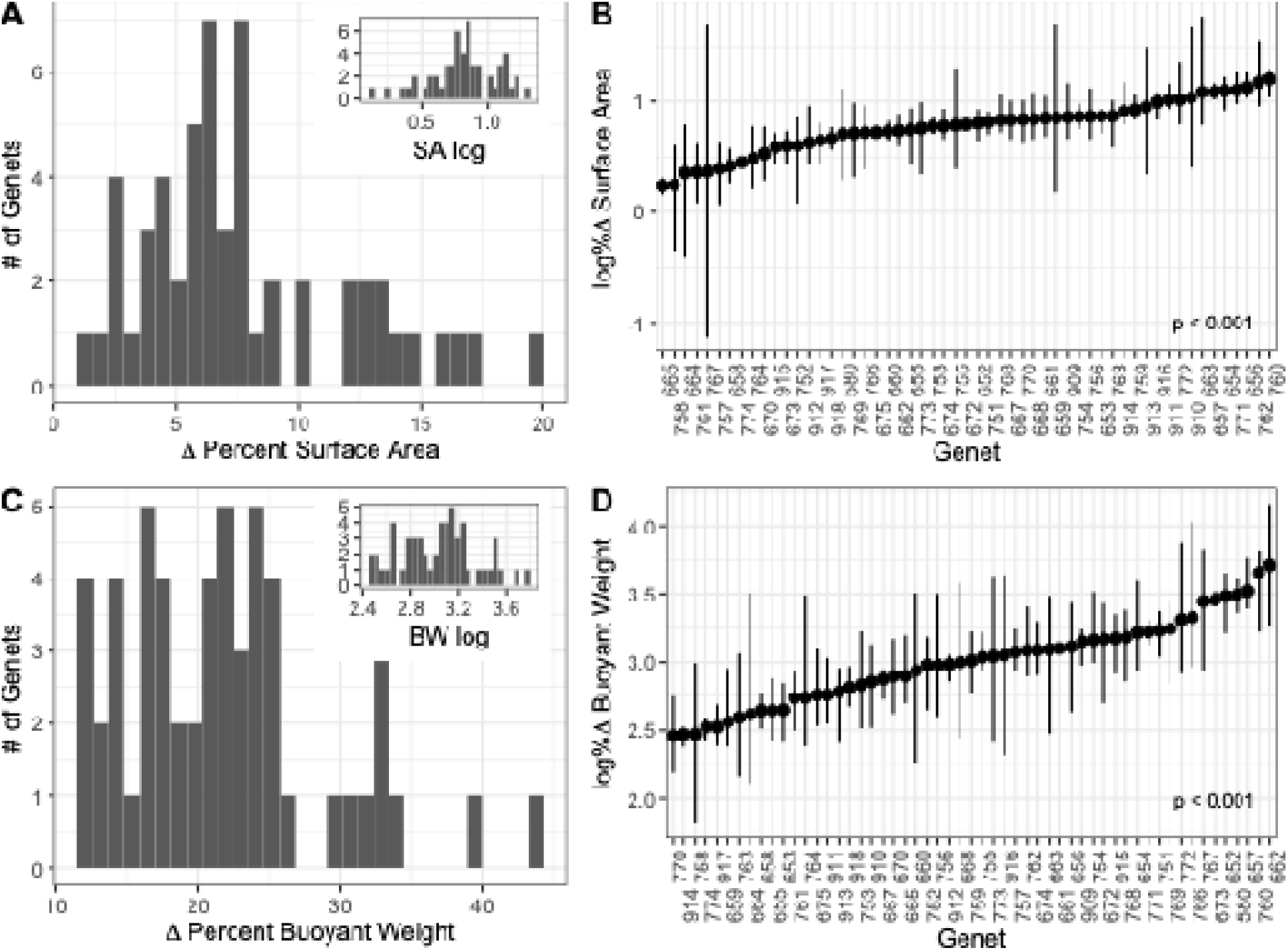
Adult growth. (A & C) Average percent change (A) surface area and (C) buoyant weight ((final-initial)/initial) of corals. The main histogram represents raw data, and the inset histogram represents log-transformed data used for analysis. (B & D) Log-transformed average percent change (B) surface area and (D) buoyant weight of genets, including the spread of all ramets. Genets that experienced total mortality (666, 669, 671, and 775) and one genet for which we could not build an ED30 thermal tolerance model (765) were excluded.

**Figure S3:**
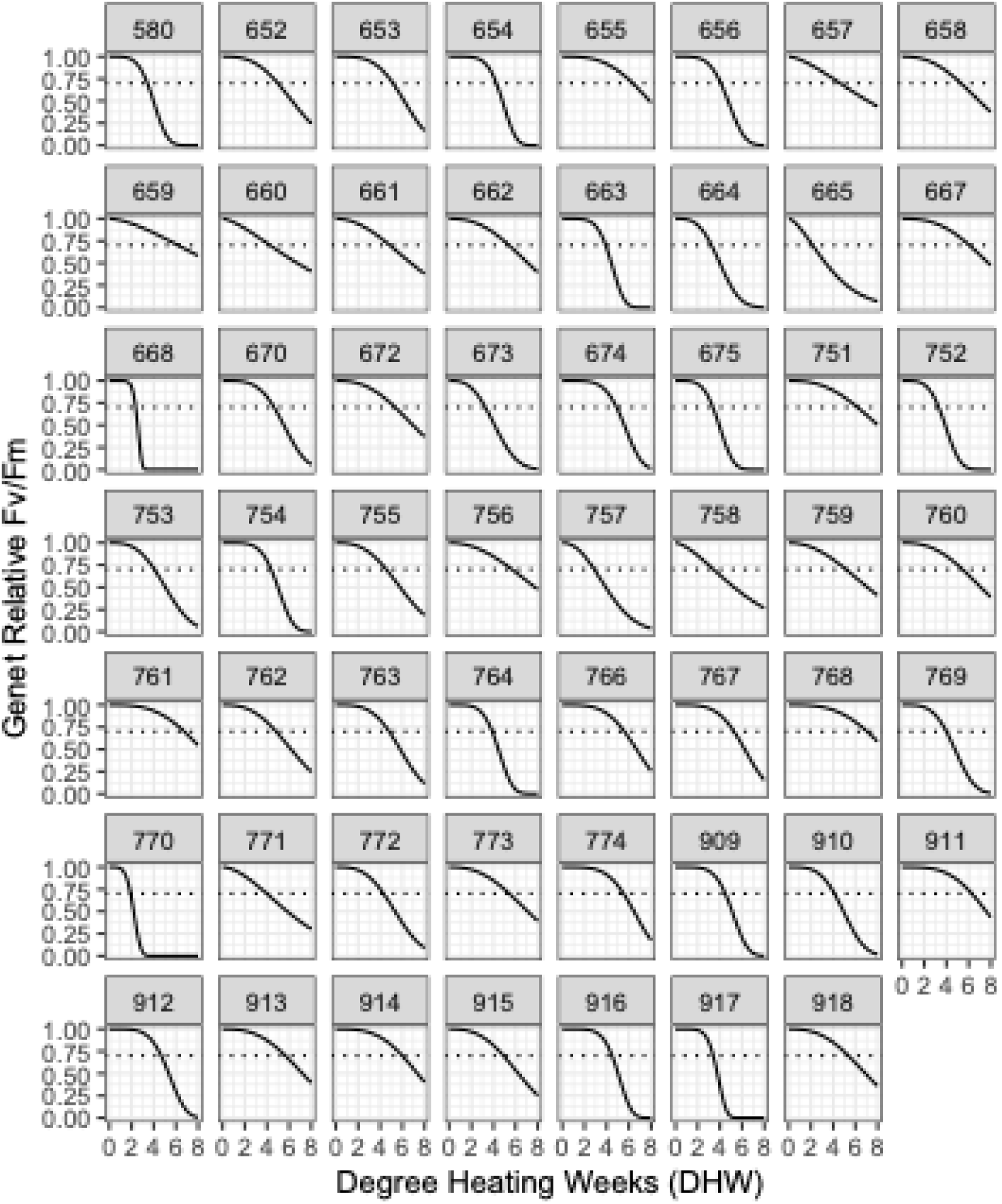
Thermal tolerance by genet. Dose response curves (ED30) based on averaged relative F_v_/F_m_ and degree heating weeks (eDHW). Thermal tolerance was defined as degree heating weeks to reach a 30% decline in F_v_/F_m_, noted by the dotted line.

**Figure S4:**
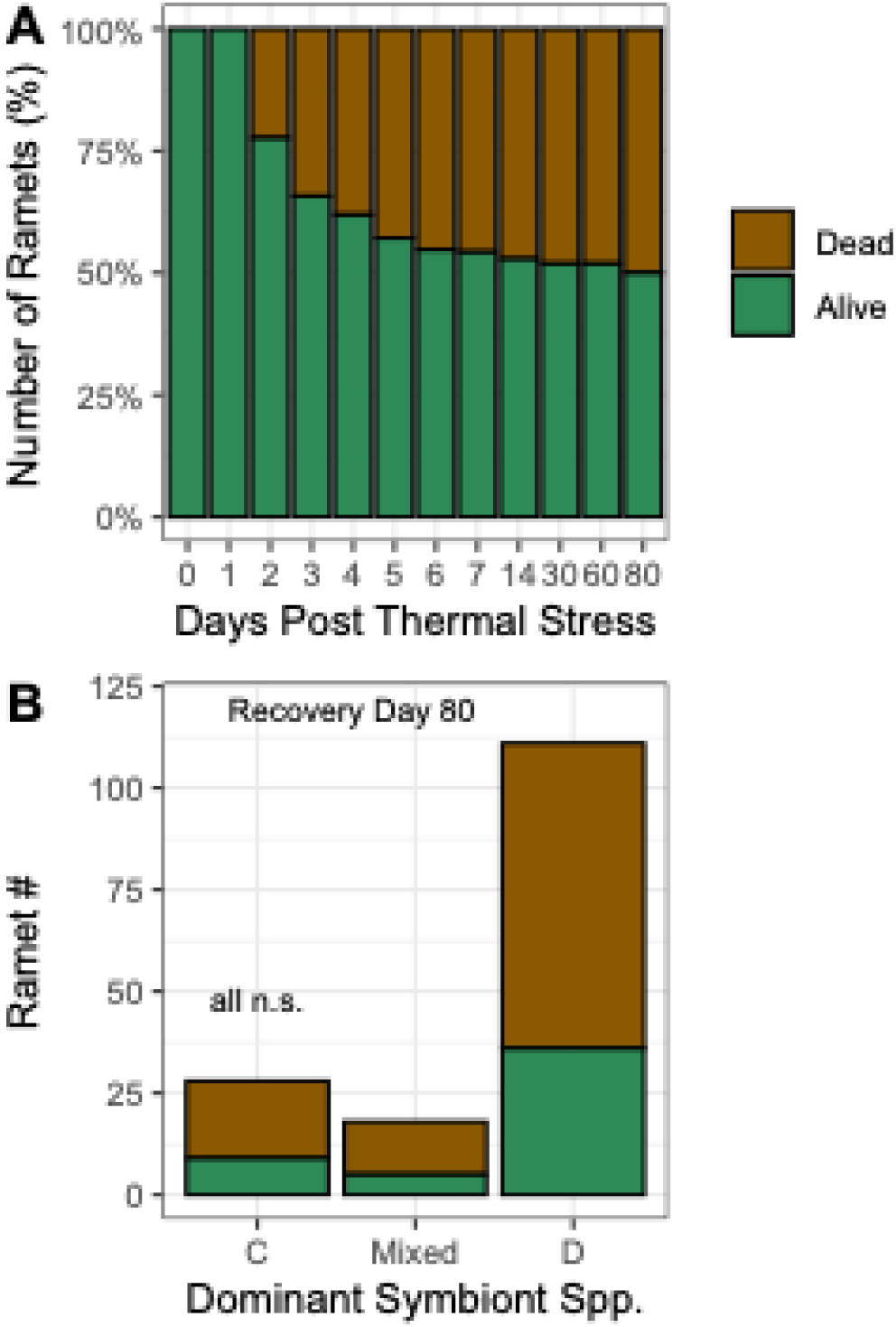
Mortality during the recovery period. (A) Ramet mortality on each sampling day during the recovery period (Days 0, 1, 2, 3, 4, 5, 6, 7, 14, 30, 60, and 80). (B) Ramet mortality at the end of recovery by symbiont community, evaluated using a fisher’s exact test, but all comparisons were not significant.

