## Supplementary figures and images for "Positive associations between growth and thermal tolerance in the reef-building coral *Montipora capitata*"

### downstream statistical analyses

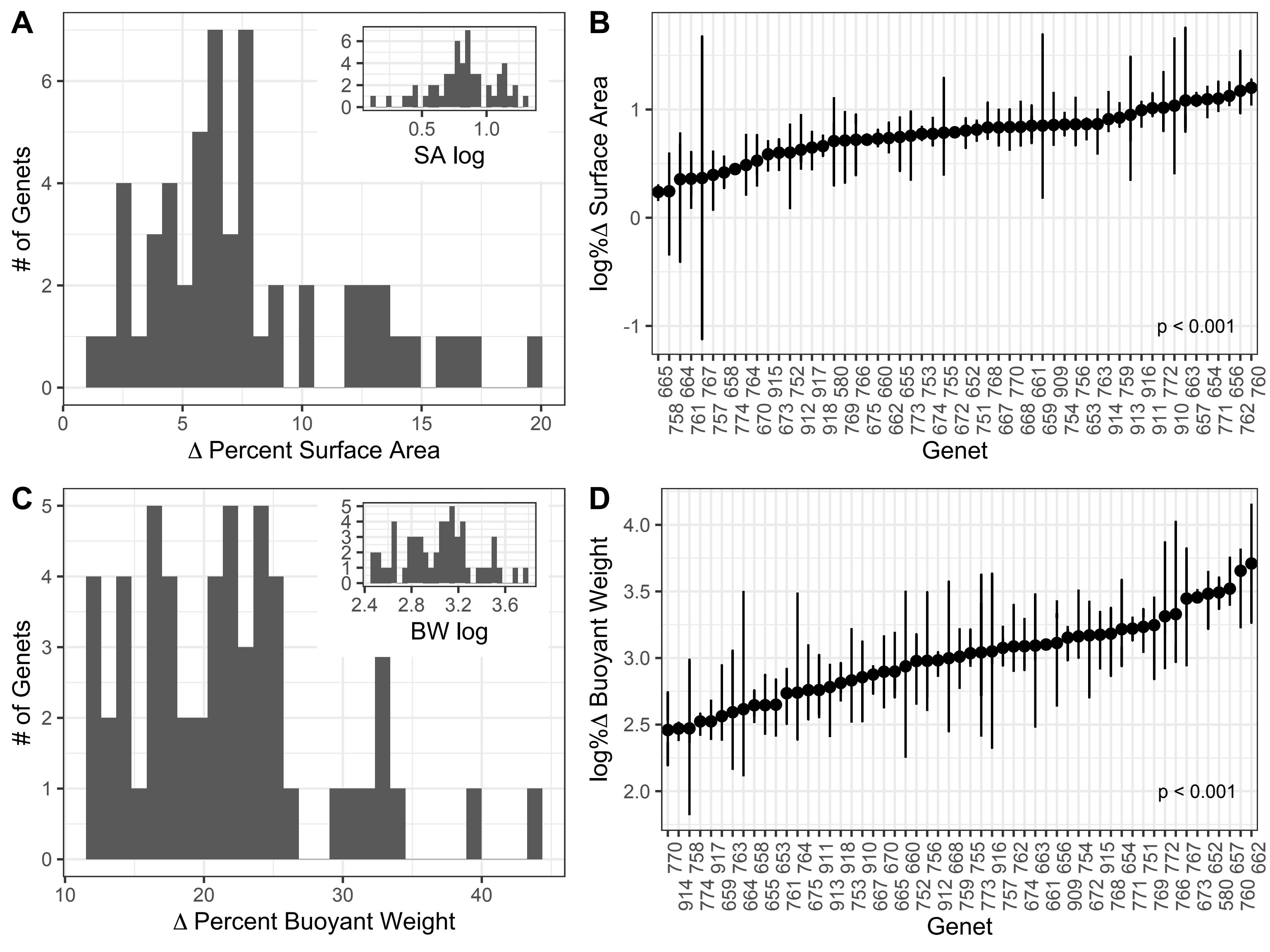

### mortality increased to 68.2

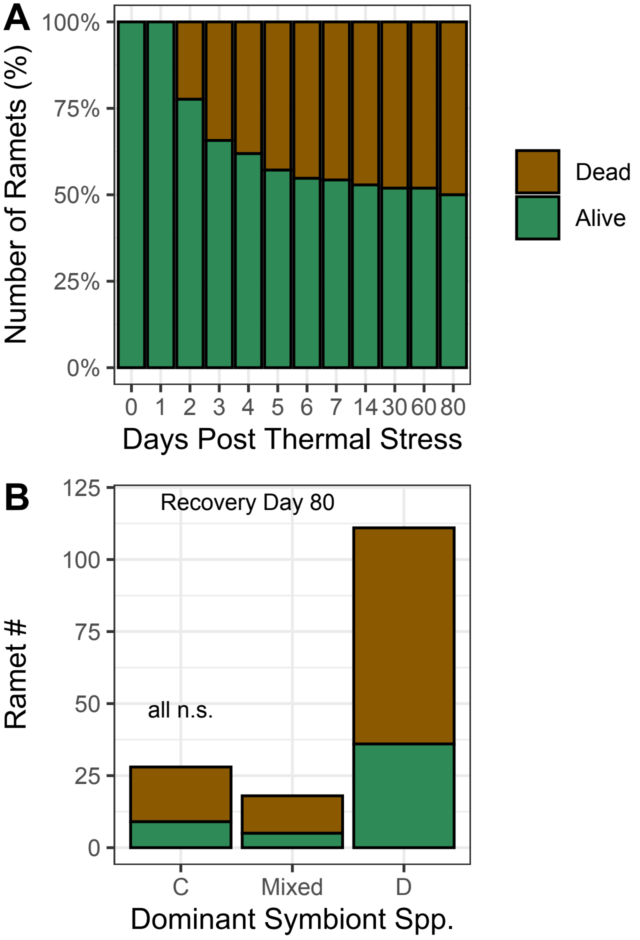

### Onset U22

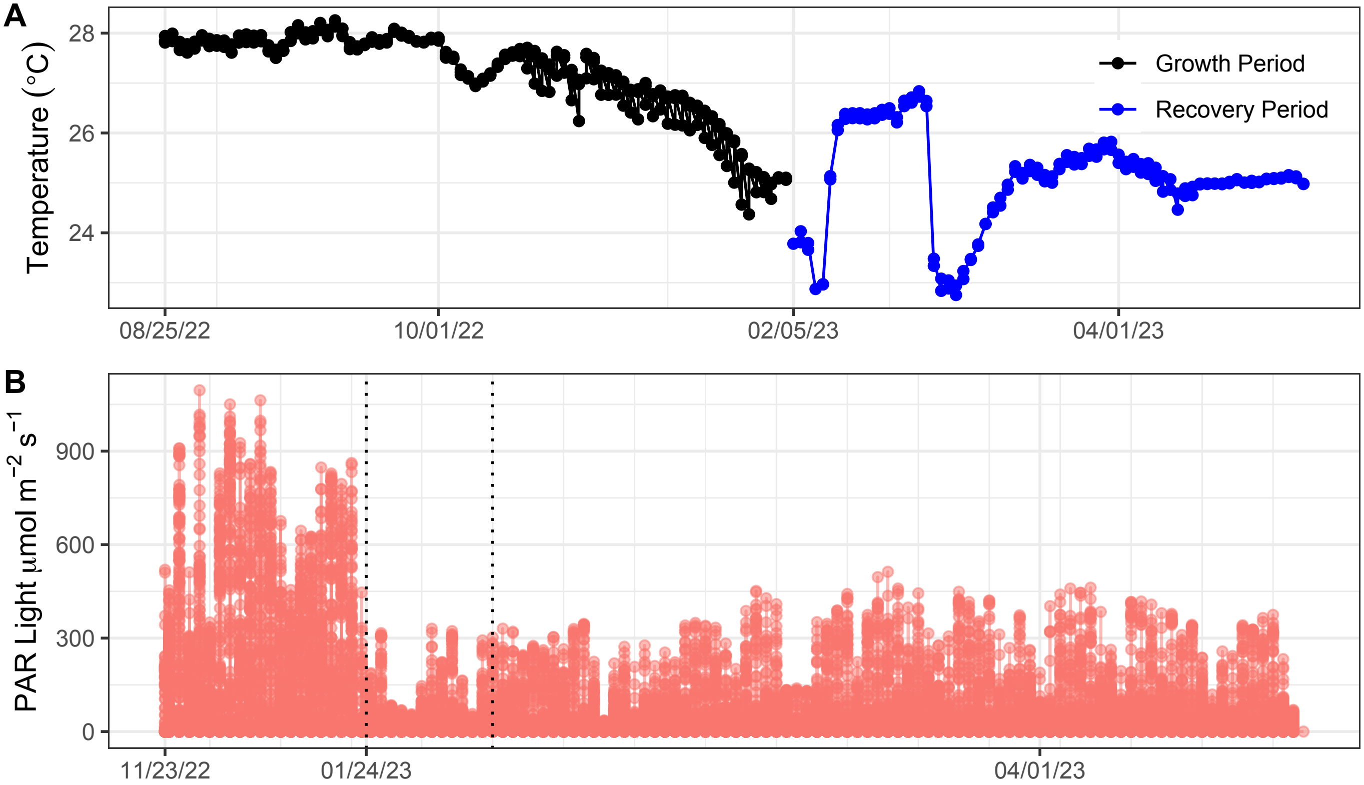

### thermal tolerance analyses

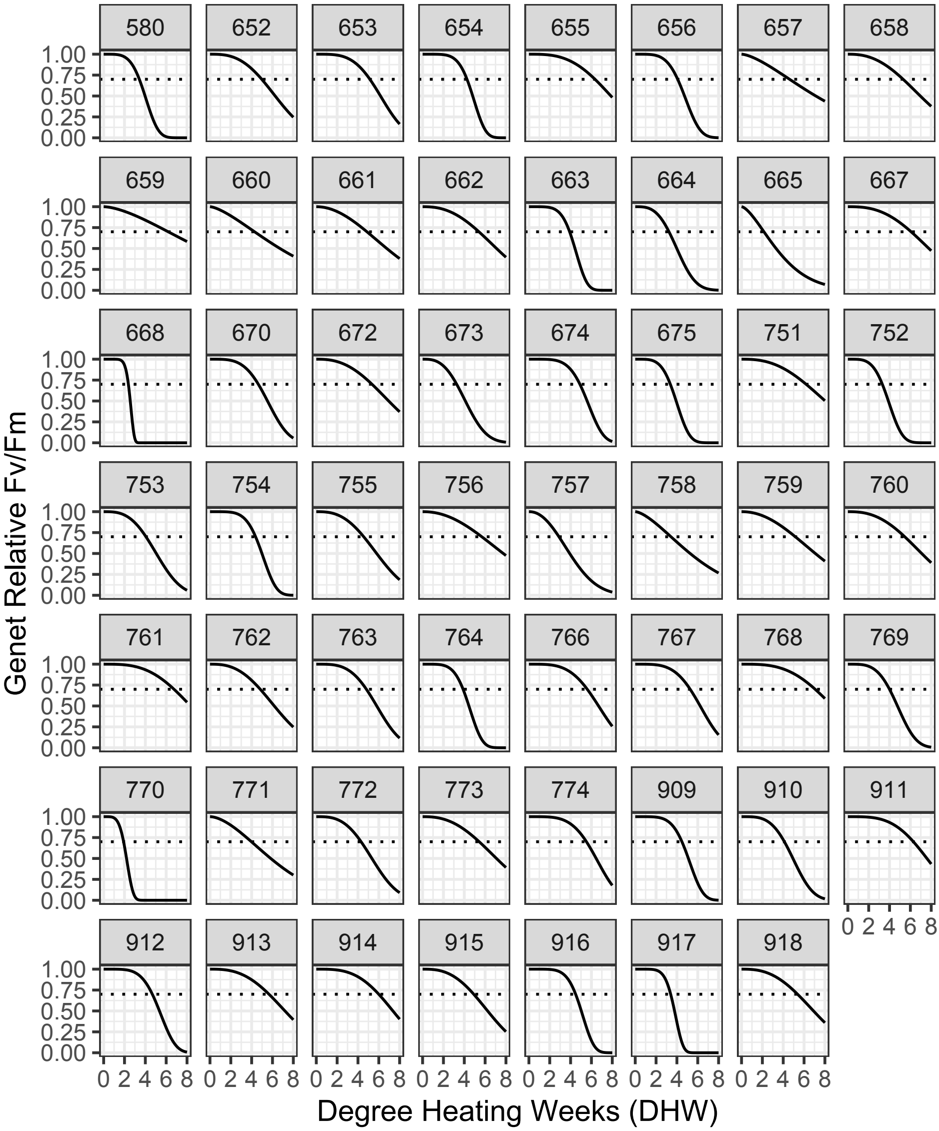
